# The fate of a mutation in a fluctuating environment

**DOI:** 10.1101/016709

**Authors:** Ivana Cvijović, Benjamin H. Good, Elizabeth R. Jerison, Michael M. Desai

**Affiliations:** Department of Organismic and Evolutionary Biology and FAS Center for Systems Biology; Department of Physics, Cambridge, MA 02138; Department of Systems Biology, Harvard Medical School, Boston, MA 02115, USA

## Abstract

Natural environments are never truly constant, but the evolutionary implications of temporally varying selection pressures remain poorly understood. Here we investigate how the fate of a new mutation in a fluctuating environment depends on the dynamics of environmental variation and on the selective pressures in each condition. We find that even when a mutation experiences many environmental epochs before fixing or going extinct, its fate is not necessarily determined by its time-averaged selective effect. Instead, environmental variability reduces the efficiency of selection across a broad parameter regime, rendering selection unable to distinguish between mutations that are substantially beneficial and substantially deleterious on average. Temporal fluctuations can also dramatically increase fixation probabilities, often making the details of these fluctuations more important than the average selection pressures acting on each new mutation. For example, mutations that result in a tradeoff between conditions but are strongly deleterious on average can nevertheless be more likely to fix than mutations that are always neutral or beneficial. These effects can have important implications for patterns of molecular evolution in variable environments, and they suggest that it may often be difficult for populations to maintain specialist traits, even when their loss leads to a decline in time-averaged fitness.

---

Evolutionary tradeoffs are widespread: adaptation to one environment often leads to costs in other conditions. For example, drug resistance mutations often carry a cost when the dosage of the drug decays [1], and seasonal variations in climate can differentially select for certain alleles in the summer or winter [2]. Similarly, laboratory adaptation to specific temperatures [3, 4] or particular nutrient sources [5, 6] often leads to declines in fitness in other conditions. Related tradeoffs apply to any specialist phenotype or regulatory system which incurs a general cost in order to confer benefits in specific environmental conditions [7]. But despite the ubiquity of these tradeoffs, it is not always easy to predict when a specialist phenotype can evolve and persist. How useful must a trait be on average in order to be maintained? How regularly does it need to be useful? How much easier is it to maintain in a larger population compared to a smaller one?

The answers to these questions depend on two major factors. First, how often do new mutations create or destroy a specialist phenotype, and what are their typical costs and benefits across environmental conditions? This is fundamentally an empirical question, which depends on the costs and benefits of the trait in question, as well as its genetic architecture (e.g. the target size for loss-of-function mutations that disable a regulatory system). In this paper, we focus instead on the second major factor: given that a particular mutation occurs, how does its long-term fate depend on its fitness in each condition and on the details of the environmental fluctuations?

To address this question, we must analyze the fixation probability of a new mutation that experiences a timevarying selection pressure. This is a classic problem in population genetics, and has been studied by a number of previous authors. The effects of temporal fluctuations are simplest to understand when the timescales of environmental and evolutionary change are very different. For example, when the environment changes more slowly than the fixation time of a typical mutation, its fate will be entirely determined by the environment in which it arose [8]. On the other hand, if environmental changes are sufficiently rapid, then the fixation probability of a mutation will be determined by its time-averaged fitness effect [9, 10]. In these extreme limits, the environment can have a profound impact on the fixation probability of a new mutation, but the fluctuations themselves play a relatively minor role. In both cases, the effects of temporal variation can be captured by defining a constant *effective* selection pressure, which averages over the environmental conditions that the mutation experiences during its lifetime. This result is the major reason why temporally varying selection pressures are neglected throughout much of population genetics, despite the fact that truly constant environments are rare.

However, this simple result is crucially dependent on the assumption that environmental changes are much slower or much faster than *all* evolutionary processes. When these timescales start to overlap, environmental fluctuations can have important qualitative implications which cannot be summarized by any effective selection pressure, even when a mutation experiences many environmental epochs over its lifetime. As we will show below, this situation is not an unusual special case, but a broad regime that becomes increasingly relevant in large populations. In this regime, the fate of each mutation depends critically on its fitness in each environment, the dynamics of environmental changes, and the population size.

Certain aspects of this process have been analyzed in earlier studies. Much of this earlier work focuses on the dynamics of a mutation in an infinite population [11–24]. However, these infinite-population approaches are fundamentally unsuitable for analyzing the fixation probabilities of mutations that are neutral or deleterious on average (and even for mutations that are beneficial on average, population sizes must often be unrealistically large for this infinite population size approximation to hold). Another class of work has focused explicitly on finite populations, but only in the case where the environment varies stochastically from one generation to the next [25–31]. Later work has extended this analysis to fluctuations on somewhat longer timescales, but this work is still restricted to the special case where selection cannot change allele frequencies significantly during an individual environmental epoch [9, 32, 33].

These studies have provided important qualitative insights into various aspects of environmental fluctuations. However, we still lack both a quantitative and conceptual understanding of more significant fluctuations, where selection in each environment can lead to measurable changes in allele frequency. This gap is particularly relevant because significant changes in allele frequency are the most clearly observable signal of variable selection in natural populations.

In this work, we analyze the fate of a new mutation that arises in an environment that fluctuates between two conditions either deterministically or stochastically on any timescale. We provide the first full analysis of the fixation probability of a mutation when evolutionary and environmental timescales are comparable and allele frequencies can change significantly in each epoch. We find that even in enormous populations, natural selection is often very inefficient at distinguishing between mutations that are beneficial and deleterious on average. In addition, substitution rates of all mutations are dramatically increased by variable selection pressures. This can lead to counterintuitive results. For instance, mutations that result in a tradeoff but are predominantly deleterious during their lifetime can be much more likely to fix than mutations that are always neutral or even beneficial. Thus it may often be difficult for populations to maintain specialist traits, even when loss of function mutations are selected against on average. This can lead to important signatures on the genetic level, e.g. in elevated rates of non-synonymous to synonymous substitutions (dN/dS) [34].

## MODEL

We consider the dynamics of a mutation that arises in a haploid population in an environment that fluctuates over time. We assume the population has constant size *N* (neglecting potential seasonal changes in the size of the population) and denote the frequency of the mutant at time *t* as *x*(*t*). In the diffusion limit, the probability density function of the frequency of the mutant, *f*(*x*, *t*), evolves according to the standard single-locus diffusion equation with a time-varying selection coefficient [35]

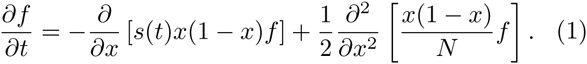

We focus on the case where the environment fluctuates between two conditions, where the (log) fitness effects of the mutation are 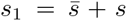 and 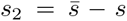, respectively. Note that 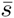 is the arithmetic average of the log fitness, which corresponds to the geometric mean of the absolute fitness. We neglect longer-term changes in selection pressures, so that *s*(*t*) will fluctuate between *s*_1_ and *s*_2_ in discrete environmental epochs (Fig. 1A). Through the bulk of our analysis we will focus on the case of a mutation with a strong pleiotropic trade-off, such that 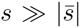 and *Ns* ≫ 1. In other words, selection in each epoch is strong compared to drift and compared to the time-averaged selection pressure. While this will not be generically true, the effects of fluctuations will turn out to be most dramatic for those mutations that fall into this regime, and we consider violations of these assumptions in the Supplementary Information. We note that this does not imply that the trait is nearly neutral on average since selection can still be strong in the traditional sense 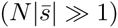.

**FIG. 1.**
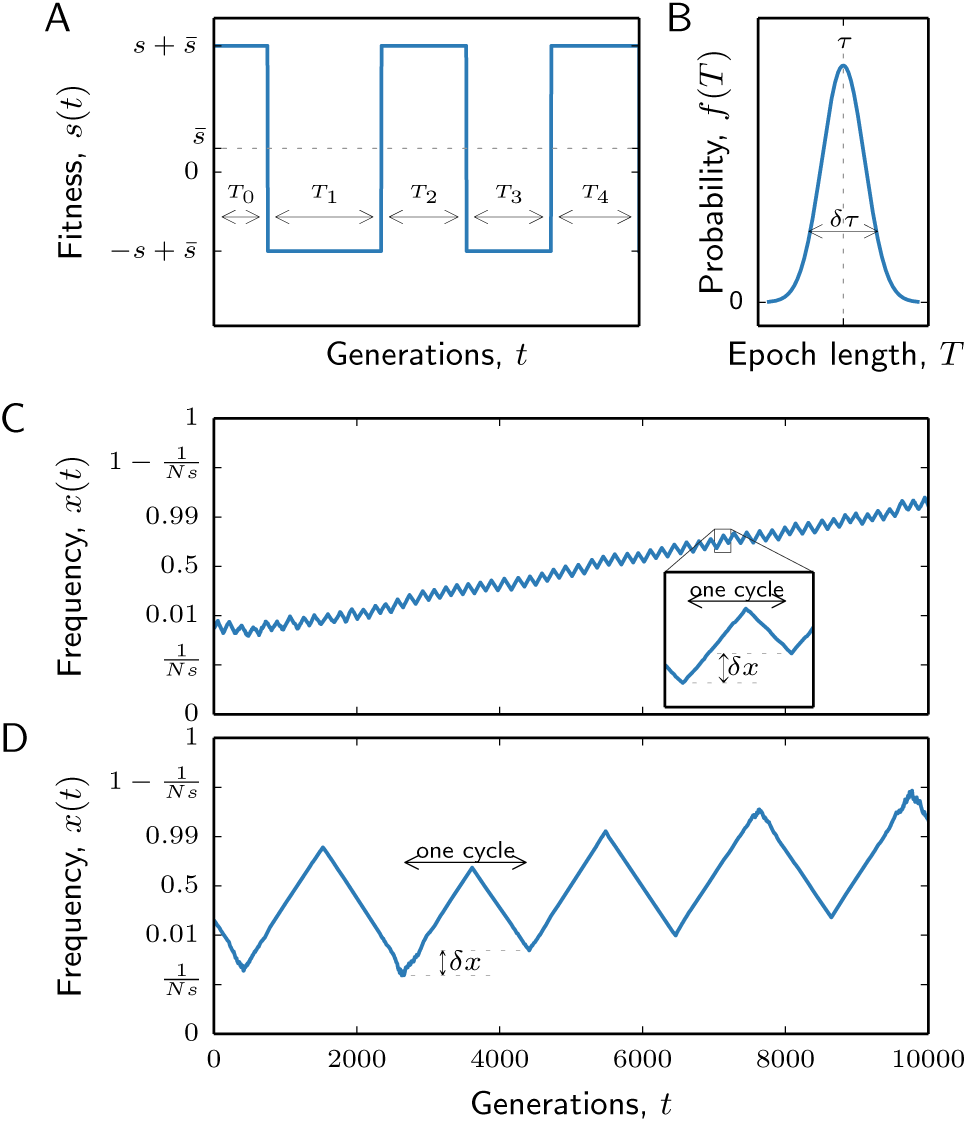
Fitness and frequency trajectories. (A) Sample fitness trajectory. The mutation arises at a random point in time. (B) Epochs have average length 〈*T*〉 = *τ* and variance var (*T*) = *δτ*^2^. (C),(D) Examples of frequency trajectories for environmental fluctuations that are (C) fast and (D) slow compared to the timescale of selection. In both panels, *N* = 10^6^, *s* = 10^−2^, *δτ* = 0.1; in panel C, 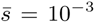, *sτ* = 1 and in panel D, 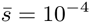, *sτ* = 10.

We assume that the duration of each epoch is drawn at random from some distribution with mean *τ* and variance *δτ*^2^ (Fig. 1B). For simplicity, we assume that the distribution of epoch lengths is the same for both environments through most of the analysis, but our approach can easily be generalized to the asymmetric case as well (see Supplementary Information). Through most of our analysis we focus on the case where the mutation rate, *μ*, is low enough that we can ignore recurrent mutation between the allelic types (*Nμ* ≪ 1). However, we show in the Supplementary Information that our analysis and conclusions also extend to the regime in which the mutation rate is high (*Nμ* ≫ 1). We discuss the relationship between our model and those employed in previous work in more detail in the Supplementary Information.

### Timescales of environmental variation

The fate of a new mutation will crucially depend on how the characteristic timescale of environmental fluctuations, *τ*, compares to the typical lifetime of a new mutation. For example, in the extreme case where environmental fluctuations are very slow, each mutant lineage will either fix or go extinct during the epoch in which it arose. Thus, its fate is effectively determined in the context of a constant environment in which it is either strongly beneficial or strongly deleterious. The fixation probability of such a mutation has been well-studied, and can be most easily understood as a balance between the competing forces of natural selection and genetic drift. We briefly review the key results here, since they will serve as the basis for the rest of our analysis below.

While the mutation is rare, genetic drift dominates over natural selection, and the mutant allele drifts in frequency approximately neutrally. When the mutation is more common, natural selection dominates over genetic drift: a beneficial mutation increases in frequency deterministically towards fixation, and a deleterious mutation declines deterministically towards extinction. To determine the threshold between these two regimes, we ask whether significant changes in allele frequency are driven by selection or drift. According to Eq. (1), natural selection changes the frequency of a rare allele substantially (i.e. by of order *x*; see [36] for details) in a time of order *t* = 1/*s* generations. In this time, genetic drift leads to a change in frequency of order 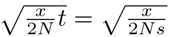. Thus there is a critical frequency 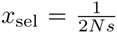 where these forces are comparable. Below *x*_sel_, genetic drift drives substantial changes in allele frequencies before natural selection has time to act, while above *x*_sel_ natural selection dominates over drift.

In the drift-dominated regime where *x* < *x*_sel_, the probability that a lineage at frequency *x* drifts to frequency *x*_sel_ before going extinct is approximately 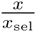.Thus a new mutation 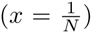 will reach this threshold with probability of order 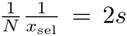. If the mutation arose during a beneficial environment, it will then grow logistically 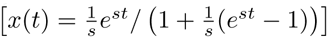 and will fix in about 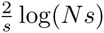 generations. On the other hand, if the mutation arose during a deleterious environment, it cannot increase in frequency substantially above *x*_sel_ and will typically go extinct within 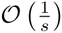 generations. Given equal probabilities of arising in either environment, the net fixation probability is therefore

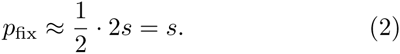

This will hold provided that the environment changes slowly enough that the mutation will have fixed or gone extinct by the end of that environmental epoch (*sτ* ≫ 2 log(*Ns*)); see Supplementary Information for further discussion and analysis of the correction due to finite epoch lengths.

In contrast, whenever *sτ* ≪ 2 log(*Ns*), a mutant lineage will experience many beneficial and deleterious epochs before it can fix. In this case, environmental fluctuations can have a dramatic influence on the frequency trajectory of a new mutation (Fig. 1). For example, when *sτ* > 1, selection within each epoch will drive the mutant frequency to very high and very low values, but because *sτ* ≪ 2 log(*Ns*), the mutation will experience many of these dramatic reversals before it fixes or goes extinct (Fig. 1D).

### An effective diffusion process

Since we aim to predict the long-term fate of the mutation, we are primarily concerned with how multiple epochs combine to generate changes in the allele frequency. This suggests that we define an *effective* diffusion process which integrates Eq. (1) over pairs of environmental epochs, similar to the earlier approaches of [32] and [9]. This yields a modified diffusion equation,

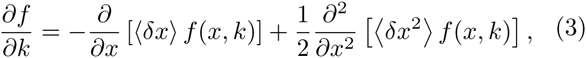

where *x* now represents the frequency of a mutation at the *beginning* of a beneficial epoch, and time is measured in pairs of epochs (Fig. 1C,D). Equation (3) also leads to a corresponding backward equation,

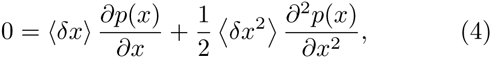

for the fixation probability, *p*(*x*), as a function of *x* [35]. Here, 〈*δx*〉 and 〈*δx*^2^〉 are the first two moments of the change in frequency in a single timestep, and must be calculated by integrating Eq. (1) over a pair of epochs. These functions will be independent of time, but will generally have a more complicated dependence on *x* than the coefficients in Eq. (1). In this way, we can reduce the general problem of a time-varying selection pressure to a time-*independent* diffusion process of a different form. The only caveat is that this process describes the fate of a mutation starting from the beginning of a beneficial epoch, while mutations will actually arise uniformly in time. Thus, we must also calculate the frequency distribution of a mutation at the beginning of its first full beneficial epoch, so that we can compute the overall fixation probability *p*_fix_ by averaging *p*(*x*) over this range of initial sizes.

In the following sections, we calculate 〈δx〉 and 〈δx^2^〉and solve the resulting diffusion equation for p_fix_ as a function of 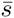, *s*, *τ*, *δτ*, and *N*. We begin by analyzing the problem at a conceptual level, to provide intuition for the more formal analysis that follows.

## HEURISTIC ANALYSIS

We first consider the simplest case of an on-average neutral mutation in a perfectly periodic environment (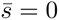, *δτ* = 0). In this case, the effects of environmental fluctuations are primarily determined by how rapidly selection acts relative to the rate of environmental change. When *τ* is much less than 1/*s*, selection barely alters the frequency of the mutation over the course of a single epoch. We can then add up the contribution of multiple epochs in a straightforward manner (see Supplementary Information), and we find that the coarse-grained process is indistinguishable from a neutral mutation in a constant environment [9, 32].

In contrast, when *τ* is much greater than 1/*s* (but still shorter than the fixation time), natural selection dramatically alters the frequency of a mutation within a single epoch, and the effects of environmental fluctuations will play a much larger role. For example, the fate of a mutation now crucially depends on the precise time at which it arises. If it arises early in a deleterious epoch, it will be driven to extinction long before the environment shifts. Since a deleterious mutation with cost *s* can survive for at most of order 1/*s* generations, the mutation must arise within the last 1/*s* generations of a deleterious epoch to avoid extinction. Similarly, if the mutation arises late in a beneficial epoch it might increase in frequency for a time, but these gains will be reversed in the subsequent deleterious epoch, when the fitness of the mutation switches to −*s* (see Figure 2A). Therefore, the mutation must arise within the first ∼1/*s* generations of a beneficial epoch in order to avoid extinction (i.e. within the “window of opportunity”, Fig. 2A). We let *τ*_*c*_ = 1/*s* denote the length of the critical period in each epoch when a successful mutation can arise. Since mutations occur uniformly throughout each epoch, only a fraction *τ*_*c*_/*τ* ≪ 1 will arise at the “right” time; all others are certainly destined for extinction.

**FIG. 2.**
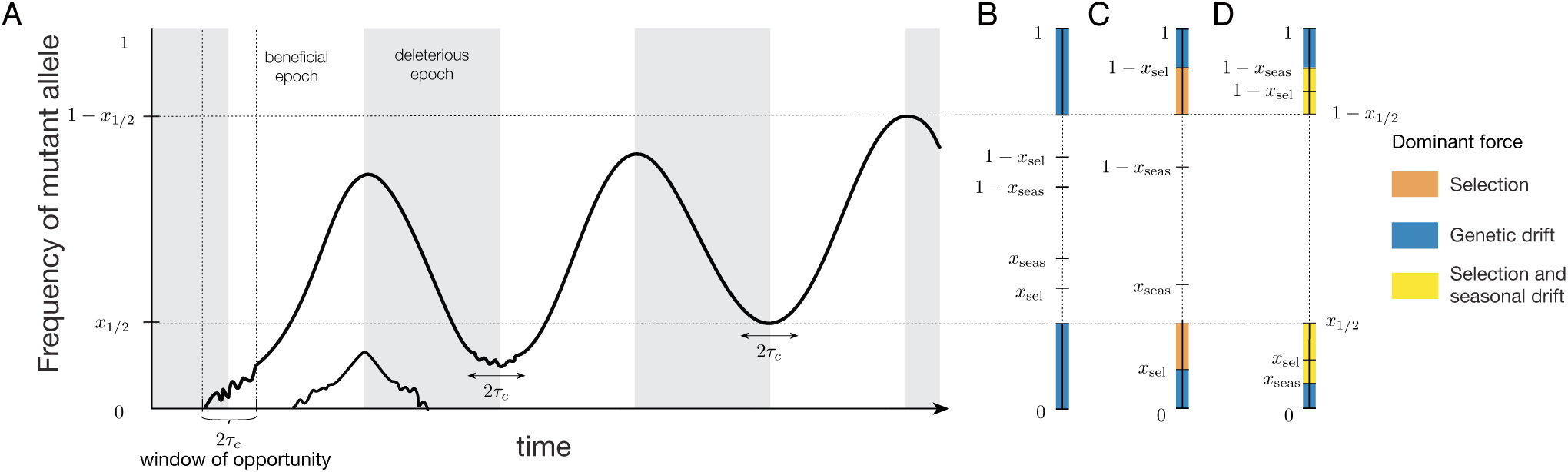
A schematic illustration of the concepts in the heuristic section. (A) All mutations that arise outside of the window of opportunity near the beginning of a beneficial epoch are destined to go extinct. Within a pair of environmental epochs, genetic drift is strongest within 2*τ*_*c*_ generations of the mutant being most rare as long as the frequency of the mutant is below *x*_1/2_ at the beginning of the beneficial epoch. If the mutant starts the beneficial epoch at *x*_1/2_, selection will take its frequency to 1 − *x*_1/2_ by the end of that epoch. The dominant evolutionary force depends on the frequency of the mutation. (B) When the average selection pressure and the variation in epoch lengths are weak, genetic drift dominates all other evolutionary forces. The mutation thus drifts neutrally below *x*_1/2_, at which point it has a fixation probability of 1/2. This picture applies regardless of whether *x*_sel_ is large or small compared to *x*_seas_. (C) When the average selection pressure is sufficiently large, *x*_sel_ ≪ *x*_1/2_ and *x*_sel_ ≪ *x*_seas_. The mutation drifts neutrally below *x*_sel_, after which its dynamics are deterministic and dominated by natural selection. This picture holds regardless of whether *x*_seas_ is large or small compared to *x*_1/2_. (D) When the variation in epoch lengths is large enough, *x*_seas_ is less than both *x*_sel_ and *x*_1/2_. The mutation first drifts neutrally below *x*_seas_. Above this critical frequency, both natural selection and seasonal drift are potentially important, depending on the magnitudes of *x*_seas_, *x*_sel_ and *x*_1/2_.

If a mutation does arise during this critical time, its future behavior is characterized by a series of dramatic oscillations in frequency, which can drive an initially rare mutant to high frequencies (and back) over the course of a single cycle (Fig. 1D). Since selection is efficient within each epoch (*Ns* ≫ 1), the effects of genetic drift are dominated by the period within of order *τ*_*c*_ = 1/*s* generations of the beginning and end of each epoch, when either the mutant or the wildtype becomes rare (Fig. 2A). However, provided that the mutation starts at a frequency *x* ≪ *e*^*-sτ*/2^, the dominant contribution to genetic drift comes from periods where the mutant is rare, since the wildtype remains above frequency *x* throughout the environmental cycle. As a result, the contributions from drift are dominated by the first *∼ τ*_*c*_ generations and the last *∼τ*_c_ generations of the cycle, when the frequency of the mutant is still close to *x*. Thus, the overall magnitude of drift is reduced by a factor of *τ*_c_/*τ*, but the dynamics of the mutation are otherwise neutral. This approximation breaks down when the frequency of the mutation is of order *e*^*-sτ*/2^, since genetic drift near the middle of the cycle (while the wildtype is rare) starts to play a larger role. This drift, when propagated to the end of the cycle, ultimately leads to a net increase in the average frequency of the mutant and the effective diffusion process is no longer neutral (see Supplementary Information).

Fortunately, by the time that the mutation reaches an initial frequency of *e*^*-sτ*/2^, we know that it must have an equal chance of fixing or going extinct. In other words, *x*_1/2_ = *e*^*-sτ*/2^ is the special frequency at which *p*(*x*_1/2_) = 1/2. This is a consequence of the inherent symmetry of the problem: when the mutant begins a beneficial epoch with frequency *x*_1/2_, the wild-type will have frequency *x*_1/2_ at the end of that epoch, and the situation will be exactly reversed – hence, the mutant and wild-type must have the same fixation probability (Fig. 2A).

Given that *p*(*x*_1/2_) = 1/2, we can calculate the fixation probability of a new mutation while it is rare, without having to consider the dynamics above *x*_1/2_. We have seen that there is a probability ∼ *τ*_*c*_/*τ* that the mutation arises at the right time; otherwise it is certain to go extinct. Provided that it arises at the right time, the mutation has an initial frequency of 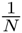, and it drifts neutrally to frequency *x*_1/2_ with 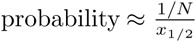 (Fig. 2B). Since it is equally likely to fix or go extinct at this point, the net fixation probability is simply

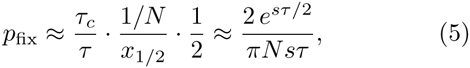

where we have also included an *𝓸*(1) factor of 4/*π*, which is derived in the formal analysis be is derived in the formal analysis below. We note that the same line of reasoning can be applied to the fast-switching (*sτ* ≪ 1) case as well, provided that we redefine *τ*_*c*_ = *τ* and *x*_1/2_ = 1/2. With these definitions, we recover the standard result that *p*_fix_ = 1/*N* when *sτ* ≪ 1 [32]. In contrast, when 1 ≪ *sτ* ≪ log(*Ns*) the fixation probability in Eq. (5) is much larger than 1/*N* (and eventually saturates to *s* when *sτ* ≫ log(*Ns*)). In other words, an on-average neutral mutation in a fluctuating environment is much more likely to fix than a strictly neutral mutation. This has important implications for the maintenance of specialist phenotypes, which we revisit in more detail in the Discussion.

### The reduced efficiency of selection

It is straightforward to extend this picture to mutations that are beneficial or deleterious on average 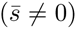. As in the constant environment case, we must consider the relative contributions of selection and drift to the net change in the mutant frequency. Over a pair of epochs, the average selection pressure will alter the frequency of the mutation by a factor of order 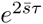, which leads to small changes of order 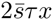 when 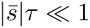. Thus, selection requires approximately 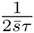 pairs of epochs to change the frequency of the mutation by of order *x*. Meanwhile, the contribution from drift over a single cycle is of order ^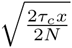^, so the cumulative drift that accumulates over 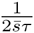 cycles is ≈ 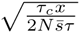. By comparing the magnitudes of these terms, we find that there is a critical frequency 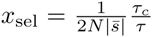 above which selection operates efficiently. If 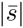 is small enough that *x*_sel_ ≫ *x*_1/2_, then the average selection pressure will not have time to influence the fate of the mutation before it reaches *x*_1/2_ (Fig. 2B), and it will fix with the same probability as Eq. (5). On the other hand, if *x*_sel_ ≪ *x*_1/2_, then the mutation will drift to frequency *x*_sel_ with 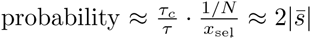, and will then deterministically fix or go extinct depending on the sign of 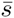 (Figure 2C). The threshold between these two behaviors occurs at 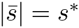, where we have defined

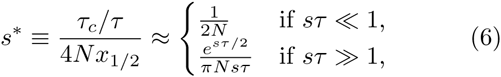

which includes an additional factor of 1/2 derived in the formal analysis below. The total fixation probability is therefore given by

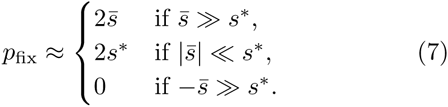

For mutations with 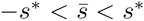, the fixation probability does not depend on the average selection coefficient and can be much higher than the fixation probability of neutral mutations in a constant environment. When fluctuations are strong (*sτ* ≫ 1), this “drift barrier” at *s*\* is much larger than the traditional value of 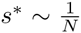 in a constant environment. Thus, we see that in addition to raising the overall fixation probability of nearly neutral mutations 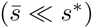, environmental fluctuations also elevate the minimum fitness effect required for selection to operate efficiently.

### The role of seasonal drift

Of course, environmental fluctuations in nature are never truly periodic, so it is natural to consider what happens when we allow for stochastic variation in the length of each epoch. To illustrate these effects, it is useful to first return to the case where 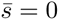. When the duration of consecutive epochs is no longer deterministic, the increase in frequency during a beneficial epoch may not always be balanced by the decrease in frequency during the following deleterious epoch. These imbalances change the frequency of the mutation by multiplicative factors of *e*^*s*Δ*T*^, which serve as an additional source of variation alongside genetic drift. However, the nature of this “seasonal drift” is very different from ordinary genetic drift, since it does not act on each individual independently. Instead, the *e*^*s*Δ*T*^ factors lead to correlated fluctuations across the whole mutant lineage. Thus, the relative changes from seasonal drift do not decrease at higher frequencies as they do for genetic drift. When *sδτ* ≪ 1, the seasonal drift over a pair of epochs leads to a change of order *sδτx*, while we have seen that the contribution from genetic drift over the same period is of order 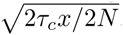. This means that there is a critical frequency 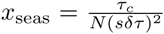 above which seasonal drift dominates over genetic drift.

If *x*_seas_ ≫ *x*_1/2_, then seasonal drift will have little time to influence the fate of the mutation before it has an equal chance of fixing or going extinct (Fig. 2B and Fig. 2C), and the fixation probability will remain the same as Eq. (5). On the other hand, if *x*_seas_ ≪ *x*_1/2_, or

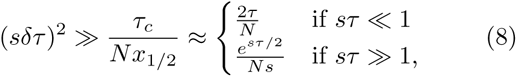

then there will be a broad range of frequencies where seasonal drift is the dominant evolutionary force (Fig. 2D). In large populations, this condition can be satisfied even when *sδτ* (and *s*τ) are extremely small. For frequencies above *x*_seas_, the multiplicative changes of seasonal drift cause the logarithm of the mutant frequency to undergo an unbiased random walk, so that the probability of reaching *x*_1/2_ before returning to *x*_seas_ is approximately log (*x*/*x*_seas_) / log *x*_1/2_/*x*_seas_). The probability that the mutation reaches the seasonal drift region (i.e. that it drifts to *c* · *x*_seas_ for some order one constant *c*) is proportional to 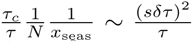. The total fixation probability is therefore of order

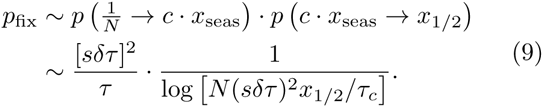

Since the right hand side of Eq. (9) is much larger than 1/*N* in this regime, we see that just a small amount of seasonal drift can dramatically enhance the fixation of on-average neutral mutations, even when *sτ* ≪ 1. In addition, since *p*_fix_ now decays as a logarithm of *N*, the relative enhancement becomes even more pronounced in larger populations.

The addition of selected mutations 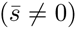 can be treated in an analogous manner, except that we must now compare the strength of selection with both genetic and seasonal drift. If 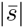 is sufficiently large that *x*_sel_ ≪ *x*_seas_, the mutation will reach frequency *x*_sel_ with probability 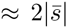 and fix or go extinct deterministically as before (regardless of whether *x*_seas_ is large or small compared to *x*_1/2_; Fig. 2C). On the other hand, when *x*_sel_ *≫ x*_seas_, selection primarily operates in the seasonal drift regime (Fig. 2D), where the logarithm of the mutation frequency undergoes a *biased* random walk with mean 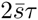 and variance (*sδτ*)^2^. When 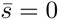, seasonal drift requires roughly log^2^(*x*_1/2_/*x*_seas_)/(*sδτ*)^2^ pairs of epochs to carry a mutation from *x*_seas_ to *x*_1/2_. If the relative change due to 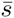 is small over this timescale, then the average selection pressure will barely bias the trajectory of the mutation before it reaches *x*_1/2_, and the fixation probability will be identical to the on-average neutral case in Eq. (9). This will be true provided 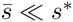, where we now have

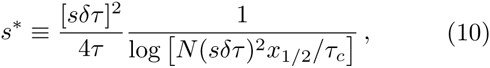

which includes the appropriate factor of 1/2 derived in the formal analysis below. On the other hand, if 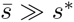, then selection dominates over seasonal drift and the fixation probability again approaches either 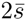 or 0. Thus, we see that seasonal fluctuations again lead to a fixation probability of the form in Eq. (7), but with *s*\* now defined by Eq. (10). In other words, seasonal drift also leads to an increase in the fitness effects required for natural selection to operate efficiently. But as we saw for the neutral fixation probability in Eq. (9), this increase is even more pronounced when seasonal drift becomes important.

## FORMAL ANALYSIS

We now turn to a formal derivation of the results described above. We begin by calculating the moments of the effective diffusion process in Eq. (4). As in the heuristic analysis above, we will work in the limit that 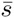*τ* ≪ 1 and *sδτ* ≪ 1. When either of these assumptions is violated, the change in frequency over a pair of epochs is no longer small and the effective diffusion approximation is no longer appropriate. We discuss violations of these assumptions in the Supplementary Information.

To calculate the moments of the effective diffusion, we must integrate the dynamics in Eq. (1) over an entire environmental cycle. When environmental switching is fast (*sτ* ≪ 1), the frequency of the mutant lineage cannot change substantially within the cycle. The overall changes in the frequency of the mutant can therefore be obtained from a short-time asymptotic expansion of Eq. (1) derived in the Supplementary Information. We can then average over the epoch lengths to obtain the moments of the effective diffusion equation

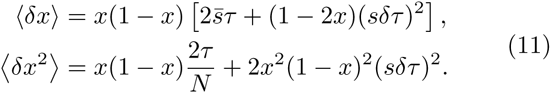

In the absence of seasonal drift (*δτ* = 0), we recover the standard moments for a mutation with fitness effect 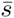 in a constant environment, where time is measured in units of 2*τ* generations. When *δt* > 0, seasonal drift leads to additional terms in both the mean and variance of *δx*, consistent with the multiplicative random walk described in the heuristic section.

These short-time asymptotics break down when environmental switching is slow (*sτ* ≫ 1), since we can no longer assume that the frequency of the mutation is approximately constant during a cycle. In this case, however, we can now model the peaks of each cycle (when either the mutant or wildtype is rare) using standard branching process methods, with asymptotic matching at intermediate frequencies. Provided that the mutant is not so common that it is likely to fix over the course of the cycle (*x* ≪ 1 − *e*^*sτ*^/*N s*), we show in the Supplementary Information that the moments of the effective diffusion equation are given by

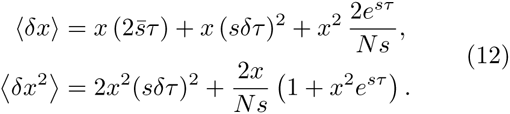

When *x* ≪ *x*_1/2_, these moments are similar to the fast-switching regime above, except that genetic drift is reduced by a factor of *τ*_*c*_/*τ* = 1/(*sτ*). For *x* ≳ *x*_1/2_, we see that additional terms arise due to genetic drift near the middle of the cycle, which increase both the mean and variance of *δx*.

In order to extend this solution to frequencies above *x* ≳ 1 − *e*^*sτ*^/*N s*, it is useful to consider the corresponding diffusion process for the wildtype frequency. By construction, the moments of this effective diffusion process are identical to Eq. (12) (with 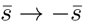)and the two sets of moments now cover the entire range of mutant frequencies. We can then find the total fixation probability *p*(*x*) by matching the corresponding solutions of Eq. (4) at some intermediate frequency where both sets of moments are valid (e.g. at *x* = *x*_1/2_). Finally, we obtain the fixation probability of a *new* mutation by averaging over the frequency of the mutant lineage at the beginning of the first full cycle it encounters. We carry out these calculations in detail in the Supplementary Information.

In both the fast and slow switching limits, we find that the fixation probability of a new mutant in a fluctuating environment satisfies a modified version of Kimura’s formula,

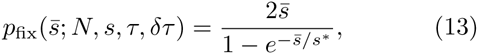

where *s*\* is defined in Eqs. (6) and (10). Equation (13) shows that the relevant fitness effect is the average fitness 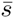, but that environmental fluctuations lead to a modified drift barrier *s*\*, which is independent of 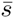 but depends on the other parameters: *N*, *s*, *τ*, and *δτ*. We compare this predicted parameter collapse to the results of Wright-Fisher simulations in Fig. 3, and compare our predictions for *s*\* with simulations in Fig. 4. These results are in full agreement with our heuristic analysis: mutations with average fitness effect 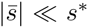 will fix with a probability approximately equal to 2*s*\*, beneficial mutations with 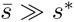 will fix with probability 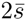, and deleterious mutations with 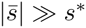 will have an exponentially small probability of fixation given by 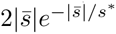.

**FIG. 3.**
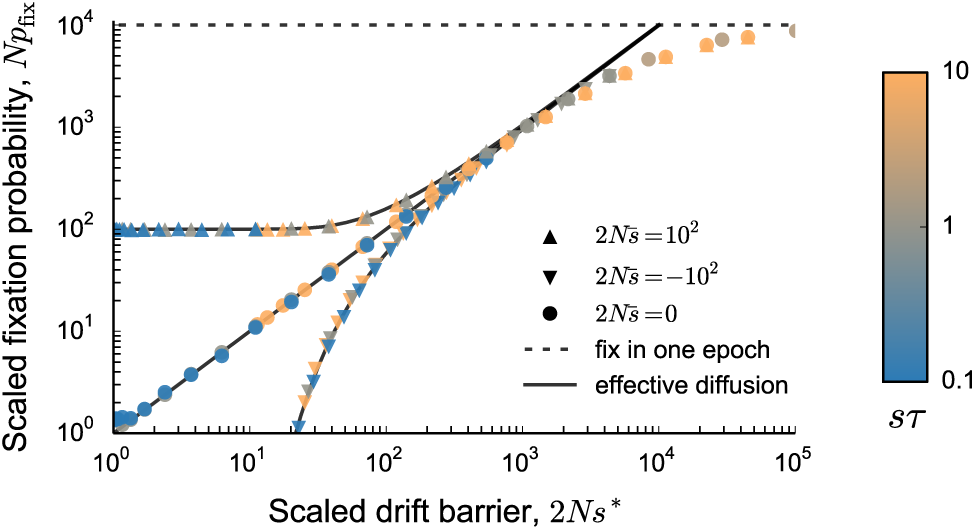
The effects of environmental fluctuations on the fate of a new mutation are well summarized by a change in the drift barrier, *s*\*. Here, *s*\* is independent of the average fitness, 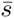, but depends on the population size and the dynamics of environmental fluctuations. Colored points show Wright-Fisher simulations of mutant lineages arising at random points in time, performed for a range of epoch lengths and variances in epoch time. Here *N* = 10^6^, *s* = 10^−2^, and var (*τ*)/*τ*^2^ varies from 10^−4^ to 10. The different colors distinguish between simulations in which switching rates were different and the different shapes distinguish between mutations that are on average beneficial (upward triangles), neutral (circles) and deleterious (downward triangles). The full lines show the theoretical predictions for the fixation probability in the effective diffusion limit (Eq. (13)) and the dotted line shows the probability of fixation in a single environmental epoch (Eq. (2)).

**FIG. 4.**
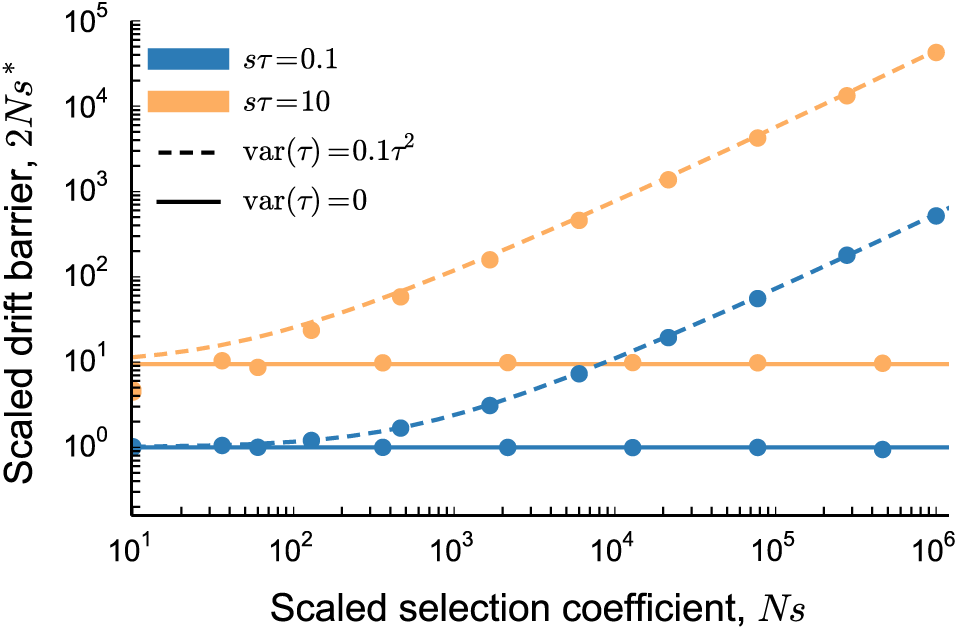
The increase in drift barrier, *s*\*, relative to its value in a constant environment as a function of the strength of selection, *Ns*. The value of *s*\* was measured using Wright-Fisher simulations of an on average neutral mutant (symbols). Lines show theoretical predictions. Fast switching (*sτ* = 0.1) is shown in blue and slow switching (*sτ* = 10) in orange. Here *s* = 10^−2^ and *N* ranges from 10^3^ to 10^8^ to obtain the values of *Ns* shown.

## DISCUSSION

In this work, we have analyzed how temporal fluctuations alter the dynamics and fixation probability of a new mutation. We find two main qualitative impacts. First, fluctuations reduce the efficiency of selection. This efficiency is commonly quantified by the ratio of fixation probabilities of beneficial and deleterious mutations, 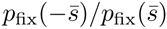. We have shown here that this ratio continues to exhibit a simple exponential dependence on 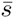,

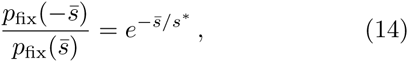

even in the presence of environmental fluctuations. As in a constant environment, Eq. (14) implies that selection cannot distinguish between beneficial and deleterious mutations when 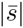 is less than the “drift barrier” *s*\*, and that selection becomes exponentially more efficient for mutations with 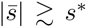. We have shown here how environmental fluctuations increase the drift barrier *s*\*, broadening the range over which selection cannot distinguish between beneficial and deleterious mutations.

Given the similarity of Eq. (14) to the constant environment case, where 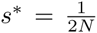, it is tempting to define an “effective population size” *N*_*e*_ = 2/*s*\*. This would attribute the decreased efficiency of selection to an increased variance in offspring number arising from variability in the environment. However, we have shown that this intuition is misleading, since the offspring number fluctuations caused by environmental variation do not affect individuals independently. This leads to behavior which cannot be captured by an effective population size [e.g., neutral fixation times which do not scale as *N*_*e*_ but rather as 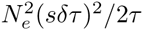.

These correlated fluctuations are also responsible for the second effect of environmental fluctuations: an overall increase in the fixation probability of all mutations. This increased rate of fixation can lead to counterintuitive results. For example, consider a mutation that is deleterious on average (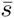 < 0) in a fluctuating environment. As is apparent from Fig. 5, the fixation probability of such a mutation can be much larger than 1/*N*, the fixation probability of a mutation that is neutral in both environments (e.g. a strictly neutral synonymous mutation). In fact, a mutation that is on average deleterious can be more likely to fix than a mutation that is on average beneficial, depending on the statistics of environmental fluctuations relevant to the two (e.g. see crossover between blue and orange lines in Fig. 5). In particular, if we compare the deleterious mutation above to a beneficial mutation of the same magnitude in a constant environment, the ratio of their fixation probabilities is given by

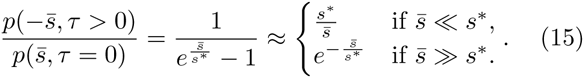

**FIG. 5.**
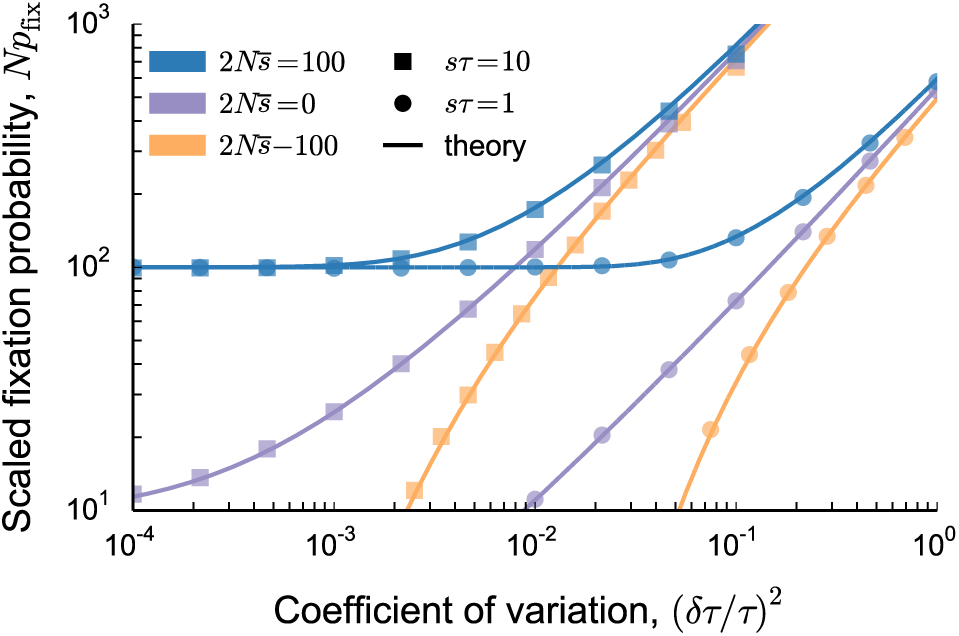
The dependence of the fixation probability on the rate and regularity of environmental fluctuations. The fixation probability has been scaled by the fixation probability of a neutral mutation in a constant environment, 1/*N*. In all simulations, *N* = 10^6^, *s* = 10^*-*2^, and the other parameters are shown in the plot. As the variance increases, the fixation probability becomes higher and the average fitness effect, 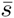, plays an increasingly smaller role. The fixation probability is also higher if the environmental changes are slower.

Due to the dramatic increase in *s\** by environmental fluctuations (Fig. 4), this ratio can often be much greater than one, reflecting a higher substitution rate of on-average deleterious mutations with a fluctuating selection coefficient compared to always beneficial mutations of the same average magnitude. The fate of a mutation can thus be more strongly influenced by the dynamics of environmental fluctuations than by its average fitness effect. At some level this is not surprising, since this behavior trivially arises whenever a deleterious mutation sweeps to fixation in a single beneficial epoch (and *p*_fix_ ≈ *s*). However, our results show that this is still true even when environmental changes are rapid enough that the mutation experiences many beneficial and deleterious epochs in its lifetime. This implies that fluctuations can accelerate sequence divergence and increase quantities such as *dN/dS* even when the population is not adapting on average. This potential consequence of fluctuating selection on rates of adaptation has been pointed out previously in the context of slow environmental fluctuations, and analyzed using the concept of “fitness flux” [10].

Our findings have important implications for the maintenance of regulatory functions in the face of a changing environment. In contrast to previous work which primarily focuses on traits which are essential in one of the two environments [7, 37], our analysis here applies to traits with more subtle costs and benefits (see [38] for a recent review). For example, bacterial regulatory mechanisms can provide an important advantage in a specific environment, but are typically costly otherwise (e.g. in the case of the *lac* operon *s* ≈ ±10% [39]). Assuming that environmental changes occur on the order of a day (*τ* ≈ 10 generations) and that *N* can easily exceed 10^6^, these populations will likely be in the regime where 1 ≲ *sτ* ≪ 2 log(*Ns*). Depending on the time spent in each environment, our analysis shows that the population can be extremely susceptible to invasion by loss-of-function mutations even if the regulatory mechanism provides an overall benefit across environmental conditions. This can make it much more difficult for a population to maintain the regulatory mechanism, leading to a “Muller’s ratchet”-like effect in which the time-averaged fitness declines over time. Furthermore, it may be equally difficult to maintain regulatory traits even in very large populations, since the drift barrier declines only logarithmically with *N* when environmental fluctuations are irregular.

**FIG. 6.**
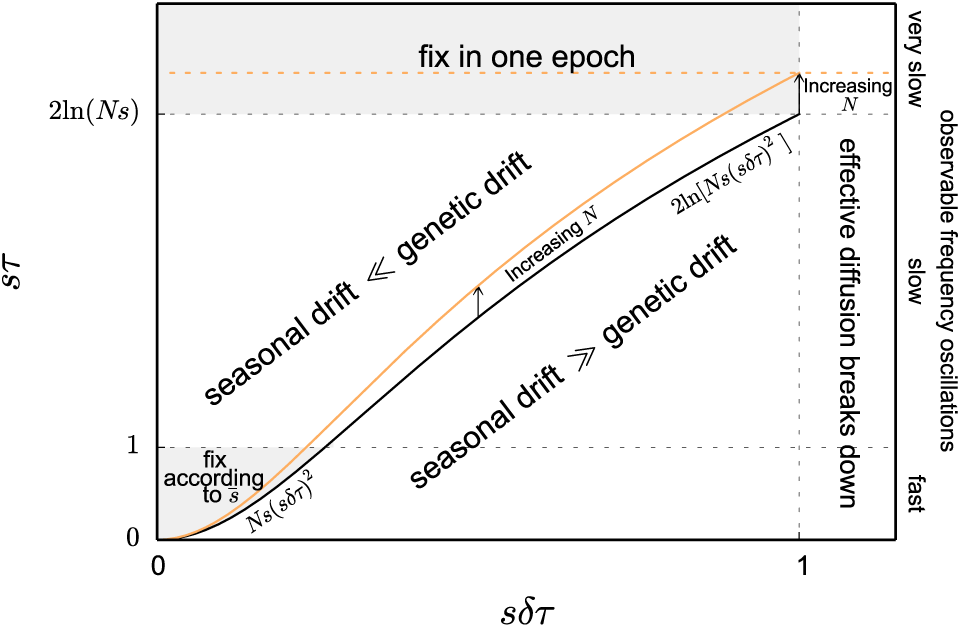
Phase diagram showing the various regimes discussed in the paper, as a function of the magnitude of environmental fluctuations (*sδτ*) and the average timescale of environmental fluctuations (*sτ*). The shaded regions are the only ones in which the environmental fluctuations do not change the drift barrier, and so the effect of environmental fluctuations can be summarized by an effective fitness. The black line separates the region in which genetic drift is the dominant source of stochastic fluctuations in the lineage size from the region in which seasonal drift has a more significant effect. The effect of an increase in the population size on the boundaries of the regions is shown in orange.

In addition to predicting fixation probabilities, our results also specify the regimes in which the evolutionary process is altered as a result of changing environmental conditions. We might have assumed that fate of a mutation is determined by its average strength of selection whenever it experiences many beneficial and deleterious epochs over the course of its lifetime (i.e. whenever *sτ* < 2 log(*Ns*)). When environmental fluctuations are both rapid and extremely regular (*sτ* ≪ 1 and 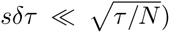 this is indeed the case. However, our analysis shows that there is also a broad regime in which environmental fluctuations lead to dramatic changes in the evolutionary process that cannot be summarized by a simple change in the effective selection coefficient (see Figure 6). This can happen for two reasons: (1) either selection within each environment is strong enough, or the duration of each epoch is long enough, that *sτ* is no longer vanshingly small, or (2) environmental fluctuations are sufficiently irregular that seasonal drift becomes important (Fig. 6).

It is not a priori clear which regime is most relevant for natural populations, largely due to the difficulty in measuring time-varying selection pressures in their native context. For a randomly chosen combination of *s* and *τ*, the rate of environmental fluctuations will often be either very fast or very slow, and the behavior described here will not apply. However, the region between these two limits becomes larger as the size of the population increases (see Figure 6), both because longer fixation times permit more extreme frequency oscillations and also because genetic drift becomes weaker relative to seasonal drift. Moreover, given a distribution of fitness effects of new mutations, it is natural to expect that some alleles will exhibit long-lived oscillations of the type studied here. Tradeoffs in this regime are arguably the most likely to be directly observed in natural populations, precisely because they exhibit frequency changes that can be measured from time-course population sequences.

For example, a recent study has identified numerous polymorphisms in natural *D. melanogaster* populations that undergo repeated oscillations in frequency over the course of the year (10 generations) [2]. Although the oscillations in many of these SNPs are likely driven by linkage to other seasonally selected sites, these data suggest that there are at least some driver alleles with *sτ* ≈ 1. The annual variation in the sizes of these populations may contribute important effects that our model does not consider, but in a population of *N* ≈ 10^5^ individuals, seasonal drift would be more significant than genetic drift as long as *δτ/τ* ≫ 0.01, corresponding to a variance in the lengths of seasons on the order of a single day.

In our analysis so far, we have primarily discussed the case where mutations incur a strong pleiotropic tradeoff and the average selection coefficient is much less than 1/*τ*. When either of these conditions is violated, the fate of a mutation is predicted by its time-averaged fitness effect and does not otherwise depend on the dynamics of environmental variation (see Supplementary Information). We have also assumed that the variance in epoch lengths is not too large, so that the changes due to seasonal drift in each cycle are small (*sδτ* ≲ 1). When this assumption is violated, the effective diffusion approximation in in Eq. (3) can technically no longer be applied. However, many of our heuristic arguments remain valid, and we expect qualitatively similar behavior of the fixation probability. We leave a more detailed treatment of this regime for future work.

## ACKNOWLEDGMENTS

We thank Eric Kang, Dmitri Petrov, Dan Rice, and Joshua Weitz for useful discussions and helpful comments on the manuscript. Simulations in this article were run on the Odyssey cluster supported by the FAS Division of Science Research Computing Group at Harvard University. This work was supported in part by the James S. McDonnell Foundation, the Alfred P. Sloan Foundation, the Harvard Milton Fund, grant PHY 1313638 from the NSF, and grant GM104239 from the NIH.

## SI Appendix

### I. THE EFFECTIVE DIFFUSION REGIME

We analyze the fate of a mutation in a fluctuating environment by employing an effective diffusion approximation, which coarse-grains the evolutionary dynamics over pairs of environmental epochs. Such an approximation is appropriate whenever the mutation experiences many beneficial and deleterious epochs over the course of its lifetime, and the net change over each cycle is small. Formally, this requires that

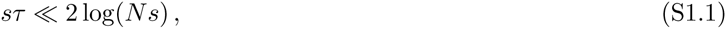

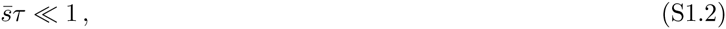

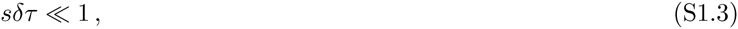

in addition to the usual strong selection assumption (*Ns* ≫ 1). We make repeated use of these limits throughout the remainder of this section. Violations of these conditions are considered in Section II.

#### A. Fast switching [*sτ* ≪ 1 ≪ 2 log(*Ns*)]

In the fast switching regime, the environmental timescale (*τ*) is much shorter than the timescale of selection (1/*s*), so the frequency of the mutation does not change much over the course of a cycle. This regime was originally analyzed by [32]; we present a derivation of these results here for completeness. We begin by rewriting the diffusion equation in Eq. [1] in Langevin form [40],

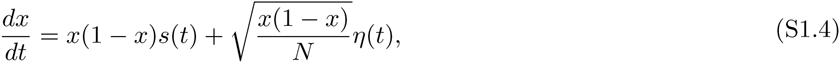

where *η*(*t*) represents the noise term and has the properties 〈*η*(*t*)〉 = 0, 〈*η*(*t*)*η*(*t*′)〉 = *δ*(*t* − *t*′). In the Itô interpretation, this can be rewritten in the following differential form

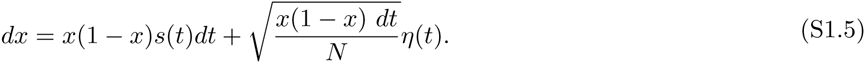

When the timescales of environmental fluctuations are shorter than the timescale of selection, 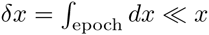, so we can assume that *x* is approximately constant over the course of a pair of epochs and coarse grain Eq. (S1.5) over an environmental cycle

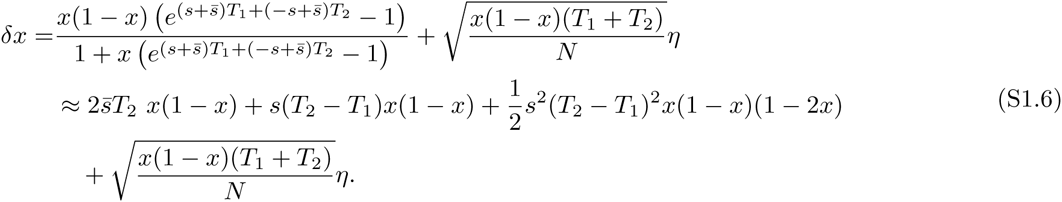

Averaging over *T*_1_ and *T*_2_, we find the first two moments of *δx*

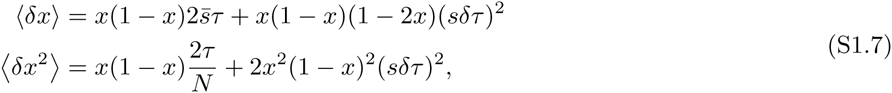

which can be rewritten as

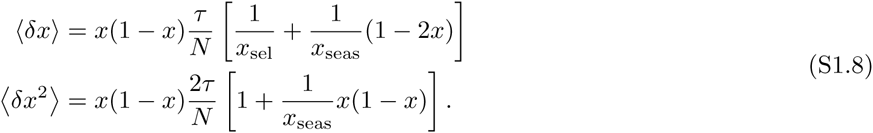

The backward equation for the fixation probability of a mutant at frequency *x* is thus [35]

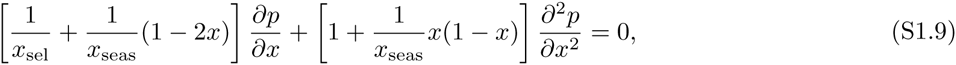

which can be rearranged as

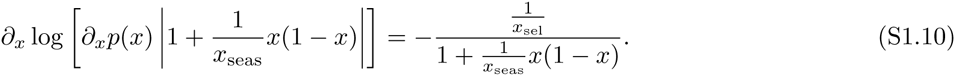

It will be convenient to define *x*_±_ to be the roots of 1 + *x*(1 − *x*)/*x*_seas_ (i.e. 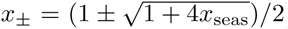). Integrating Eq. (S1.10), we obtain

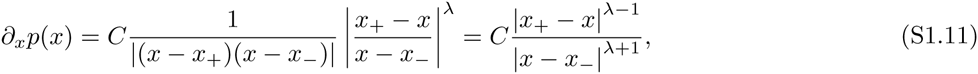

where we have defined 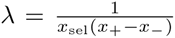. Finally, integrating Eq. (S1.11) and requiring that *p*(0) = 0 and *p*(1) = 1 gives

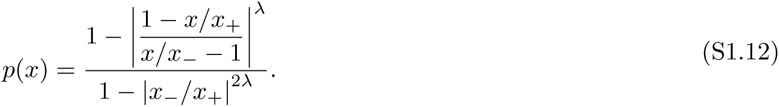

Since *sτ* ≪ 1, the initial size of the lineage will not be much greater than 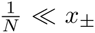 at leading order. Thus, we can expand the numerator of Eq. (S1.12) for small *x* to arrive at

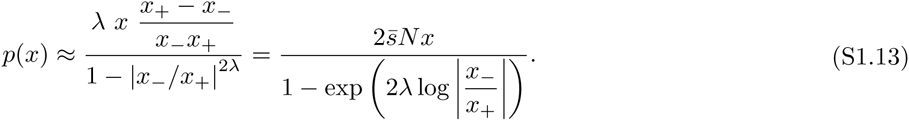

The fixation probability of a new mutation is therefore given by

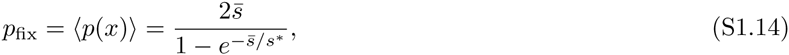

where we have used the fact that 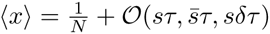, and defined the drift barrier, *s\**, as

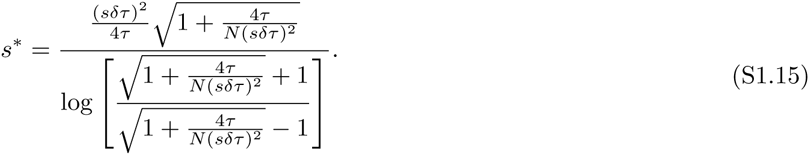

In the two limiting cases, this formula reduces to

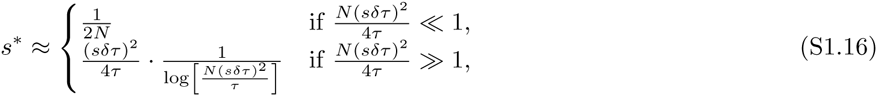

which agrees with the expressions for *s\** given in the main text.

#### B. Slow switching [1 ≪ *sτ* ≪ 2 log(*Ns*)]

In contrast to the fast switching regime above, slower environmental switching (*sτ* ≫ 1) can lead to substantial changes in allele frequency over the course of a single epoch. However, provided that 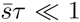 and *sδτ* ≪ 1, the net change in frequency after a full cycle is still sufficiently small. Note that when *sτ* ≫ 1, these two conditions also imply that 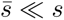 and *δτ* ≪ *τ*. In other words, our analysis simplifies to that of a *nearly* perfect fitness tradeoff in a *nearly* deterministic environment (although as we will see below, the residual effects of 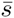 and *δτ* will still be extremely important).

To account for the nonlinear effects of selection over the course of a cycle, we begin by introducing the change of variable *χ* = *x*/(1 − *x*) in Eq. [1], which transforms the original diffusion equation into the form

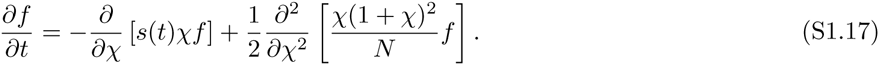

The drift term is important only at very high and very low frequencies (corresponding to *χ* ≪ (*Ns*)^−1^ and *χ* ≫ *Ns*), so we introduce a negligible error at frequencies for which *χ* ≪ *Ns* by ignoring the nonlinear component in the drift term. This gives

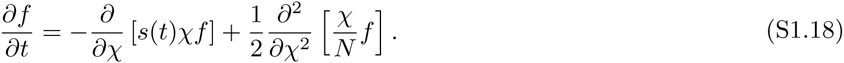

We can derive an analogous equation that is valid whenever the frequency of the wildtype is not too high by the change of variable *χ′* = 1/(1 − *x*)

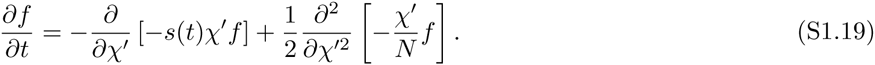

Over the course of a pair of epochs, the frequency of a mutation takes both low and high values, but we can account for the change in frequency over the entire cycle by using Eq. (S1.18) when *χ* ≤ 1 and Eq. (S1.19) when *χ′* ≤ 1 and matching the two processes at *χ* = *χ′* = 1, where they are both valid.

Concretely, let *χ* = *χ*_0_ ≪ 1 at the beginning of a beneficial epoch of length *T*_1_. We would like to calculate the moments of *δ*_*χ*_, the total change in *χ* by the end of the following deleterious epoch, which has length *T*_2_. The moment generating function of *χ*(*t*), defined as *H*_*χ*_(*z, t*) = 〈exp(-*zχ*(*t*))〉, conditioned on *χ*(0) = *χ*_0_, for an arbitrary *s*(*t*) is given by [11]

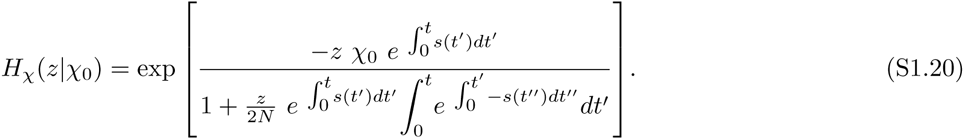

Thus, at some time *t* after the beginning of the beneficial epoch, such that 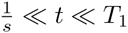, the generating function of *χ* is

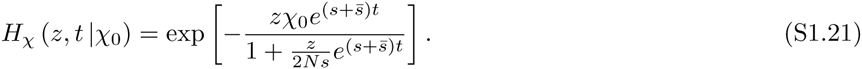

It will be convenient to define the random variable *υ*_1_ as 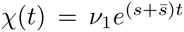. Note that *υ*_1_ captures all the non-deterministic changes in *χ*. The generating function of *υ*_1_ is

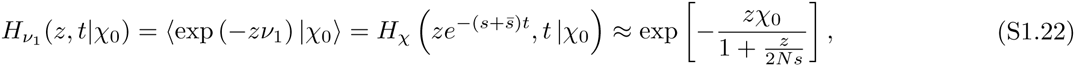

and its mean and variance are

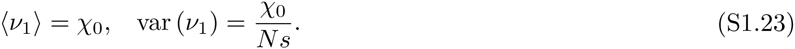

The mutation reaches *χ* = 1 at some random time 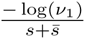, or 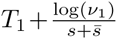 generations before the end of the beneficial epoch. From this moment on, the wildtype is the rare allele and we switch over to diffusion in *χ*′. Analogously to *υ*_1_, we define a second random variable *υ*_2_ that satisfies 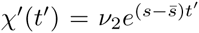, where *t*′ is the time measured from the beginning of the *deleterious* epoch from the perspective of the mutation (i.e. the middle of the cycle). Subject to the initial condition that *χ*′ = 1 at 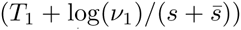 generations before the environmental shift, the generating function for *υ*_2_ at some time *t*′, such that 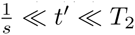 is

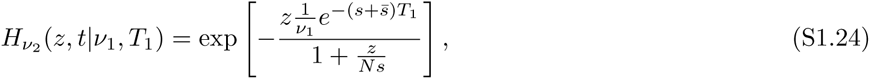

from which we obtain the conditional mean and variance of *υ*_2_

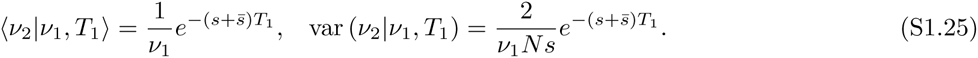

Finally, we compute the generating function for *χ* at the end of the deleterious epoch, conditioned on it having initial value 1, 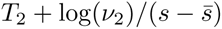 generations before the end of the deleterious epoch. We find

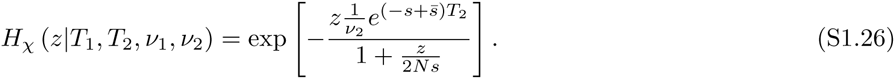

The generating function of *δχ* can be obtained from Eq. (S1.26) by noting that *H*_*δ χ*_ (*z, t*) = *e*^*zχ0*^*H*_*χ*_(*z, t*), which yields the conditional moments of *δχ*

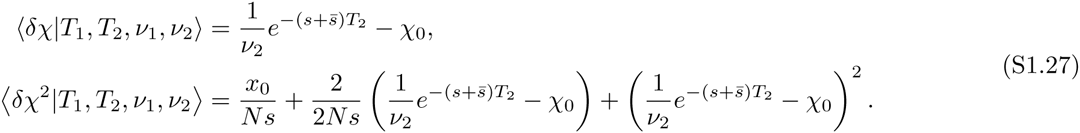

The unconditional moments are obtained by averaging over *υ*_1_, *υ*_2_, *T*_1_, and *T*_2_. In doing this we note that *T*_1_, *T*_2_ and *υ*_1_ are independent, and that *T*_2_ and *υ*_2_ are independent. We make use of the fact that the the standard deviations of all variables are much smaller than their means as long as 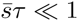 and *sτ* ≪ 1. To lowest order

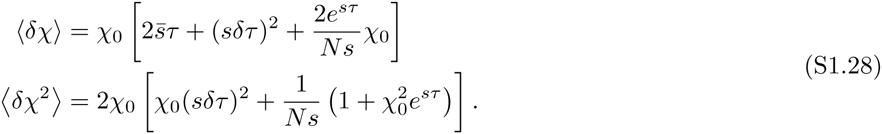

When *x* ≪ 1, *χ* ≈ *x*, so the moments of *δx* in this limit are

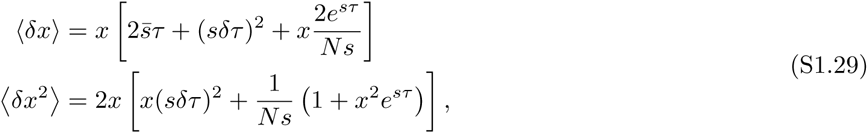

which we rewrite as

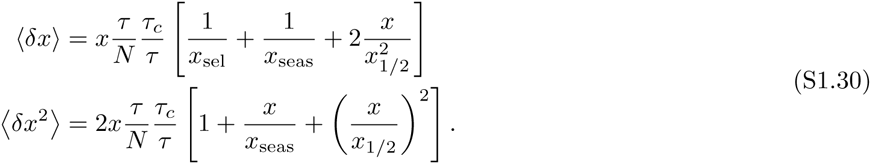

The expressions for the moments of the effective diffusion in the slow and fast switching limits (given by Eq. (S1.30) and Eq. (S1.30)) are equivalent up to the term proportional to 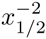. This term arises from the amplification of the effects of drift in the middle of the environmental cycle and is thus negligible in the fast switching limit. To solve the backward equation and obtain an expression for *p*(*x*), we proceed analogously to Appendix I A. Defining *x*_±_ as the roots of the polynomial 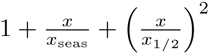 and 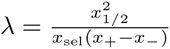, we arrive at

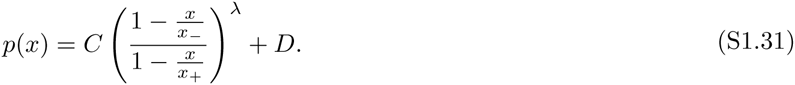

Applying the boundary condition *p*(0) = 0 and requiring that at *x*_1/2_ the probability of fixation of the mutation (and its derivative) is continuous with the probability of extinction of the wildtype at the same frequency, we arrive at

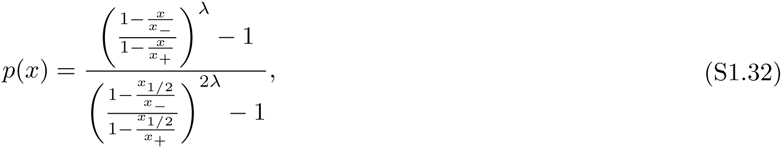

which reduces to

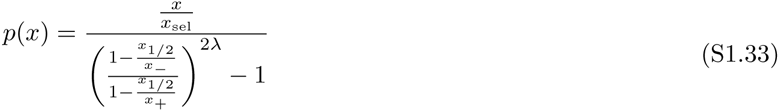

when *x* ≪ 1.

To find the probability of fixation of a new mutation arising at an arbitrary point in time, we must again average over the possible frequencies at the beginning of the first deleterious epoch. To leading order in 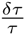, the distribution of epoch lengths is approximately *δ*-distributed,

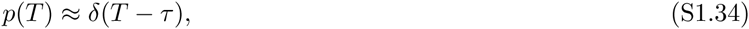

The primary contribution to the variation in initial frequencies is thus given by the random arising time, which we can average over to find that

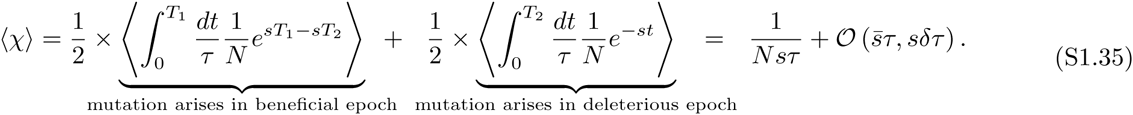

The fixation probability of a new mutation is therefore

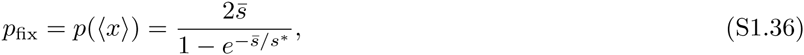

where we have defined the drift barrier,

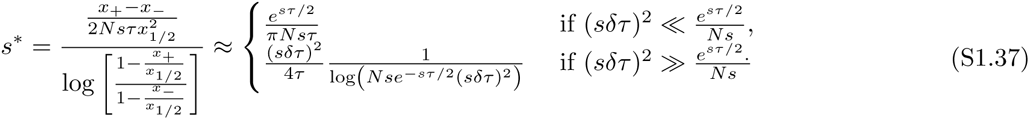

#### C. Unequal epochs (*τ*_1_ ≠ *τ*_2_)

The preceding analysis was carried out under the implicit assumption that the distribution of time spent in each environment is equal. We can relax this assumption simply by redefining the variables *s*, 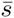, *τ* and *δτ*. For a general combination of *s*_1_, *s*_2_, *τ*_1_, *τ*_2_, *δτ*_1_, and *δτ*_2_, we can define

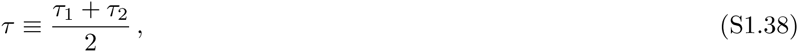

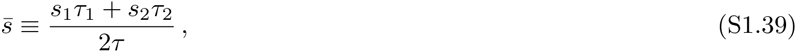

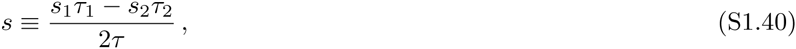

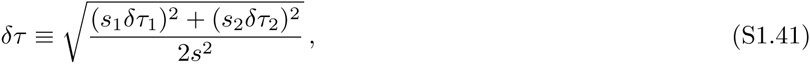

and all of our results continue to apply.

#### D. Recurrent mutation (*Nμ* ≫ 1)

In large populations, new mutations that either create or destroy a specialist phenotype might arise multiple times during the course of evolution. In this section, we consider the scenario in which wild-type individuals recurrently mutate with per-generation probability *μ* and reverse mutations from the mutant to the wild-type allelic state occur with rate *υ*. These mutation rates can encompass any mechanism by which individuals change allelic state (e.g. in prokaryotes they can include both mutations and trait gain and loss due to horizontal gene transfer). For conciseness of presentation, we will limit ourselves to the special case where *μ* = *υ*, but the analysis can be straightforwardly extended to the case where the two mutation rates are different. As before, we assume that selection within each of the environmental epochs is efficient (*Ns* ≫ 1) and stronger than mutation (*μ* ≪ *s*). When the latter is not true, selection has a very limited impact on all timescales.

In the limit that *Nμ* ⟶ 0, the entire mutant lineage will usually descend from a single mutational event and lineages that arise from different mutational events will not co-segregate in the population. Our analysis in the main text describes this regime. In contrast, when *Nμ* ≫ 1, new mutations to both of the allelic types may occur within a fixation time, and our analysis must be modified. In this recurrent mutation regime, neither of the allelic types will fix, but there will be an equilibrium distribution of frequencies at which the mutant is present in the population. Thus, instead of comparing the fixation probabilities of the two alleles, we can ask whether the average frequency of the on-average beneficial allele will be higher than the frequency of the on-average deleterious allele.

When the mutant allele is rare, genetic drift takes of order 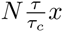 generations to change the frequency of the mutation by *x*. During this time mutation changes the allele frequency by 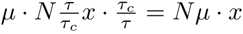, where the τ_c_/τ factor comes from the fact that the mutation must arise within the window of opportunity in order to remain in the population. Thus, when *Nμ* ≫ 1, mutation is stronger than genetic drift at all frequencies. During this time, selection will change the frequency of the mutation by 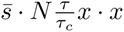 and seasonal drift will change the frequency of the mutation by 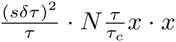 and so there will again be two critical frequencies 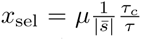 and 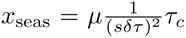, above which selection and environmental drift dominate over mutation. Genetic drift is subdominant in all of these regimes.

In the absence of seasonal drift, recurrent mutation from the wild type to the mutant will act to increase the frequency of the mutant individuals below *x*_sel_ (at these frequencies, reversion to the wild type can be neglected). Above *x*_sel_, selection will be the dominant force. If the mutation is deleterious on average, selection will decrease the number of mutant individuals above *x*_sel_ and the allele will be maintaned at equilibrium frequency *x*_sel_. As long as *x*_sel_ < *x*_1/2_, the frequency of the mutant allele averaged over time will be lower than 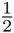. Similarly, if the mutation is beneficial on average, the frequency of the mutant individuals will be sustained at 1 – *x*_sel_ by the balance between selection and mutation from the mutant to the wild type allele. In this case, the time-averaged frequency of the mutant allele will be above 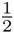 as long as *x*_sel_ < *x*_½_. When *x*_sel_ ≳ *x*_½_, selection is subdominant over the entire effective diffusion range, and mutation will sustain the time-averaged frequency of the mutant at 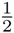. This sets a threshold for the average fitness effect at 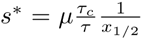 (note the similarity between this expression and the one we had in the absence of recurrent mutation, 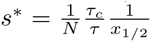). In summary, if 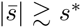, selection will be efficient at maintaining the on-average beneficial allele at a frequency that is higher than 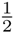. Otherwise, the average frequency of both of the alleles will be sustained at 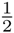.

In the presence of seasonal drift, mutational pressure will lead to the increase of the frequency of mutant alleles below *x*_seas_ and to the decrease of the frequency of mutant alleles above 1–*x*_seas_. Between *x*_seas_ and *x*_1/2_ and between 1–*x*_seas_ and 1–*x*_1/2_, selection and seasonal drift will be the dominant forces. In this case, we must compare the timescales on which selection and seasonal drift operate to determine whether or not selection is efficient. Repeating the calculation from the heuristic section in the main text, we find that selection is efficient if 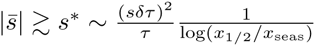.

We have seen that the same heuristic analysis applies in the *Nμ* ≫ 1 regime as the analysis we performed for *Nμ* ≪ 1 in the main text. Provided we replace *N* with 1/*μ*, we obtain analogous expressions for *s*\* and recover the same qualitative results on the efficiency of selection in a fluctuating environment. In order to make this claim more quantitative, we now turn to calculating the equilibrium distribution of frequencies for the effective process. The probability that the mutant allele frequency is below *x*_seas_ or above 1–*x*_seas_ will be suppressed by mutation. In the regime in which seasonal drift and selection dominate, the equilibrium distribution of frequencies for the effective process satisfies the equation [35]

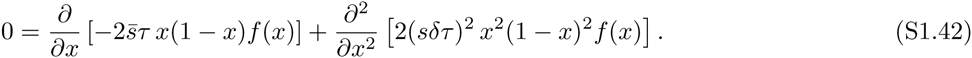

Transforming the variables to 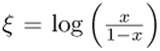, imposing zero flux boundary conditions at equilibrium, and requiring continuity of the *f*(ξ) at 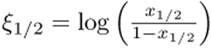 and −ξ_1/2_, we find that

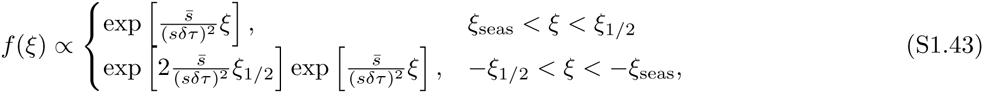

This means that *f(x)*

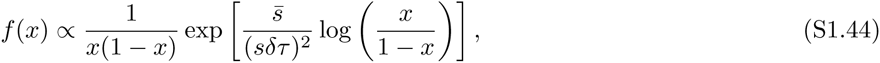

will have two peaks at *x*_seas_ and 1 − *x*_seas_. This expression for *f(x)* agrees with similar expressions in other parameter regimes that are quoted in the existing literature [26, 31, 32]. The ratio of the heights of these peaks at *x*_seas_ and 1 − *x*_seas_ is

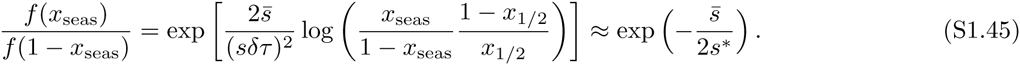

From this distribution one may in principle calculate any statistic of the frequency of the mutant allele. For instance, we use it calculate the expectation value of 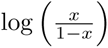, averaged over all times. This statistic will be zero if the average frequency of the mutated allele is 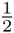, positive if the mutated allele is dominant for the majority of the time, and negative if the wild type allele is dominant in the population.

Over the course of a single environmental cycle of length *2T* starting from the beginning of a beneficial epoch

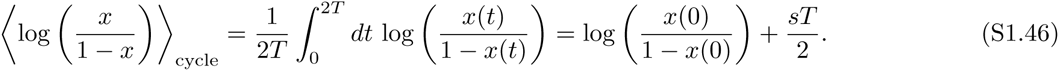

Averaged over many cycles,

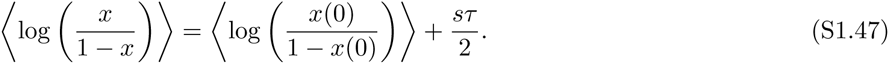

Using the analogous expression for 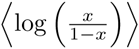 conditioned on the cycle starting in a deleterious epoch, we can calculate the expectation over the full frequency range

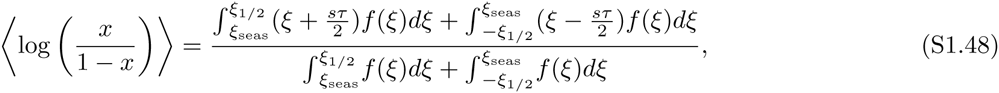

which evaluates to

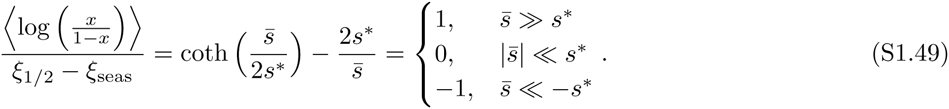

Thus, the average frequency of both of the alleles will be around 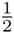 when the average selection coefficient is significantly smaller in magnitude than *s*\*, whereas the on-average beneficial allele will dominate if 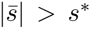, reproducing our heuristic conclusions. Of course, in this regime the effects of clonal interference across multiple loci may also become important; this is an interesting avenue for future work.

### II BEYOND THE EFFECTIVE DIFFUSION REGIME

The effective diffusion approach in the previous section relied on three basic assumptions:

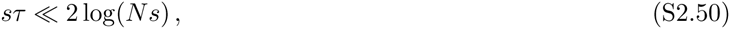

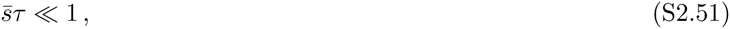

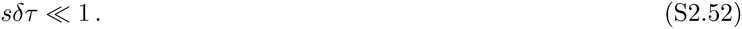

In this section, we consider violations of each of these assumptions in turn.

#### A. **Fixation during the first epoch** [1 ≪ 2 log(*Ns*) ≪ *s*τ]

The primary assumption of the effective diffusion approach is that the mutation will experience many beneficial and deleterious epochs during its lifetime. This assumption will obviously break down in the limit of extremely slow environmental switching [*s*τ ≫ 2 log(*Ns*) ≫ 1], when mutations typically fix within a single beneficial epoch. To calculate the fixation probability in this regime, we recall that to leading order in 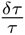, the length of the first epoch is approximately *δ*-distributed,

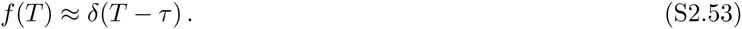

Meanwhile, the fixation time in the beneficial environment is also approximately *δ*-distributed to leading order in log(*Ns*)^−1^:

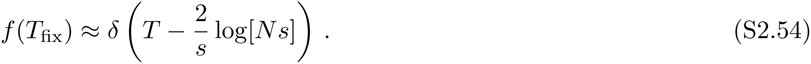

**FIG. S1.**
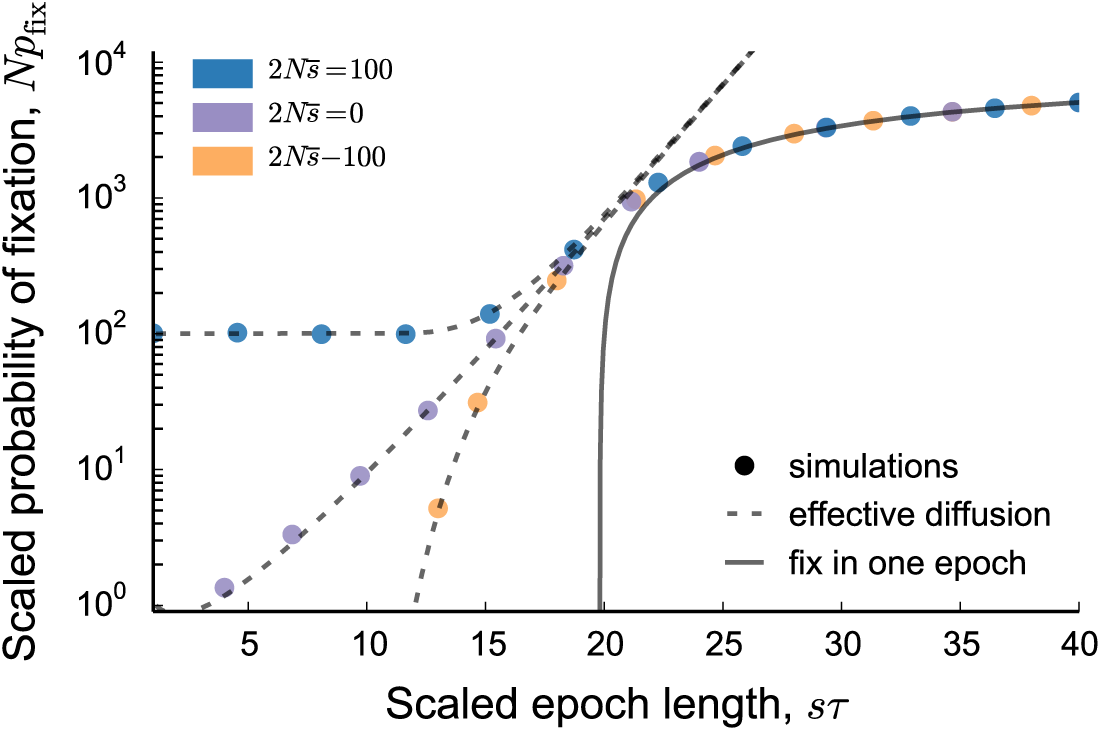
The scaled fixation probability as a function of *sτ*, which shows the transition between the effective diffusion and single epoch limits. Symbols show Wright-Fisher simulations and lines show theoretical predictions (*N* = 10^6^, *s* = 10^−2^, *δτ* = 0, and the other variables are indicated on the graph).

Thus, the primary source of variability in whether the mutation fixes stems from the random arising time of the mutation. In other words, to leading order, the fixation probability of a new mutation in this limit is given by

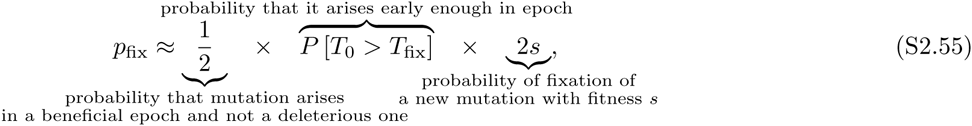

where we have used *T*_0_ to denote the time from arising to the end of a beneficial epoch. Given our assumptions above, we can evaluate the probability P [*T*_0_ > *T*_fix_] to obtain

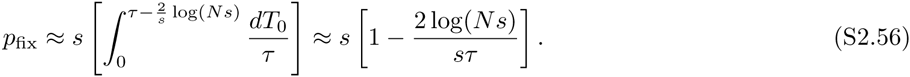

The first-order correction shows that the fix-in-one-epoch behavior breaks down when *s*τ ≲ 2 log(*Ns*), which is precisely where the effective diffusion approximation starts apply. This is illustrated in Figure S1, where we compare these predictions to simulations over a broad range of *sτ*.

#### B. **Substantial average fitness effects** (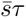 ≳ 1)

When 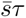 is not small compared to one, the net change over a cycle can be too large for the effective diffusion approximation to apply. In addition, when 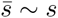, the differences between *s*_1_ and *s* can start to become important as well.

We first note that *p*_fix_ must be a monotonic function of 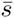, provided that we hold the remaining parameters constant. Since 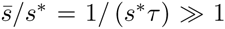, Eq. [13] shows that the fixation probability of a deleterious mutation is bounded by an arbitrarily small number, while the fixation probability of a beneficial mutation is at least 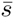. In this case, the fate of the mutation is determined while it is rare (*x* ≪ 1), which suggests that we can calculate the fixation probability by employing a linear approximation to Eq. [1],

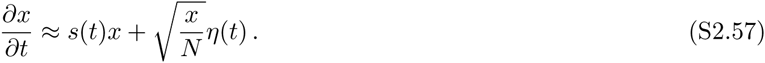

The fixation probability of this process has been well studied [11, 24], and is given by the general formula

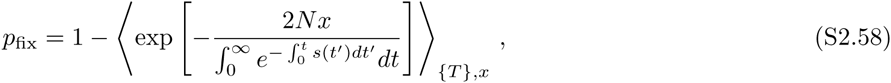

where the angle brackets denote an average over all epoch lengths as well as over the frequency of the mutation at the beginning of the first good epoch. Let *B* denote the (random) value of the integral in the denominator:

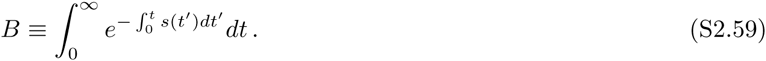

Then we can derive the following recursion relation for the distribution of *B*:

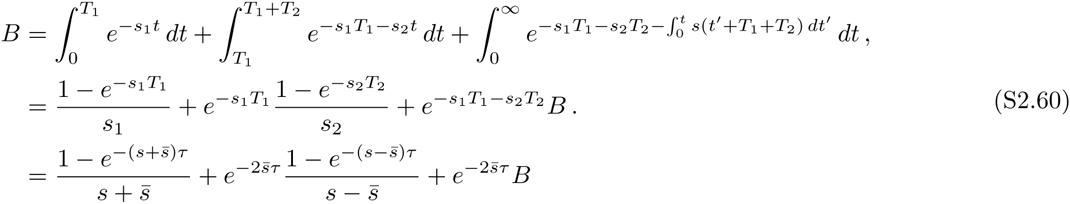

where we have retained only the leading order terms in *sδτ* and (*sτ*) ^1^. This recursion shows that to leading order, the distribution of *B* is essentially deterministic, so that we can simply solve the equation for *B* to obtain

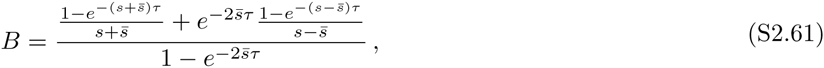

and hence

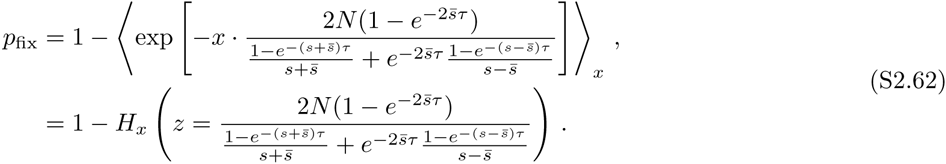

Here, *H*_*x*_(*z*) denotes the generating function of the mutant frequency at the beginning of its first full beneficial epoch. To leading order in *sδτ*, this generating function is the same as that derived in Section I B. Integrating over the possible arising times of the mutation, we find that

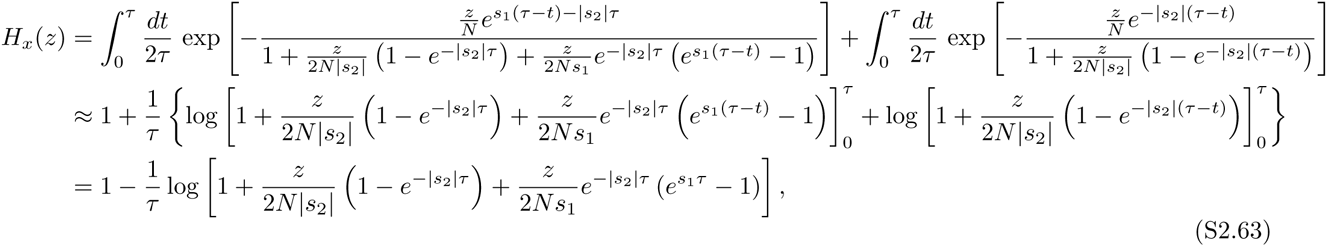

where the expansion of the exponentials is valid provided that *z* < ∞. Thus,

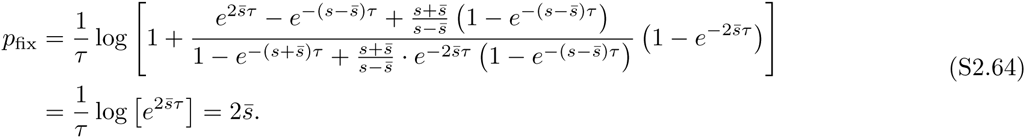

We compare this prediction with simulations in Figure S2.

#### C. **Substantial variation in epoch lengths** (*sδτ* ≫ 1)

When *sδτ* ≫ 1, seasonal drift can generate large changes in allele frequency over the course of a cycle that render the effective diffusion approximation invalid. In addition, when *δτ* ∼ *τ*, the precise shape of the epoch length distribution starts to play a larger role, and the mean and variance may not adequately capture the behavior.

However, given that the log-transformed allele frequency is still diffusive even when *sδτ* ≫ 1, we might expect our existing expressions to provide a reasonable approximation to the fixation probability, provided that the distribution of epoch lengths is still well-summarized by the mean and variance. To check this conjecture, we compare our original theoretical predictions to simulations in the slow switching regime under an exponential distribution of epoch lengths (where *sδτ* = *sτ* ≫ 1) in Figure S2. We see that our existing expressions provide a reasonable approximation to the fixation probability even when *sδτ* ≫ 1, although some small deviations are noticeable due to the modified dynamics during the first few epochs (i.e., before the mutation reaches the edge of the seasonal drift-dominated region).

**FIG. S2.**
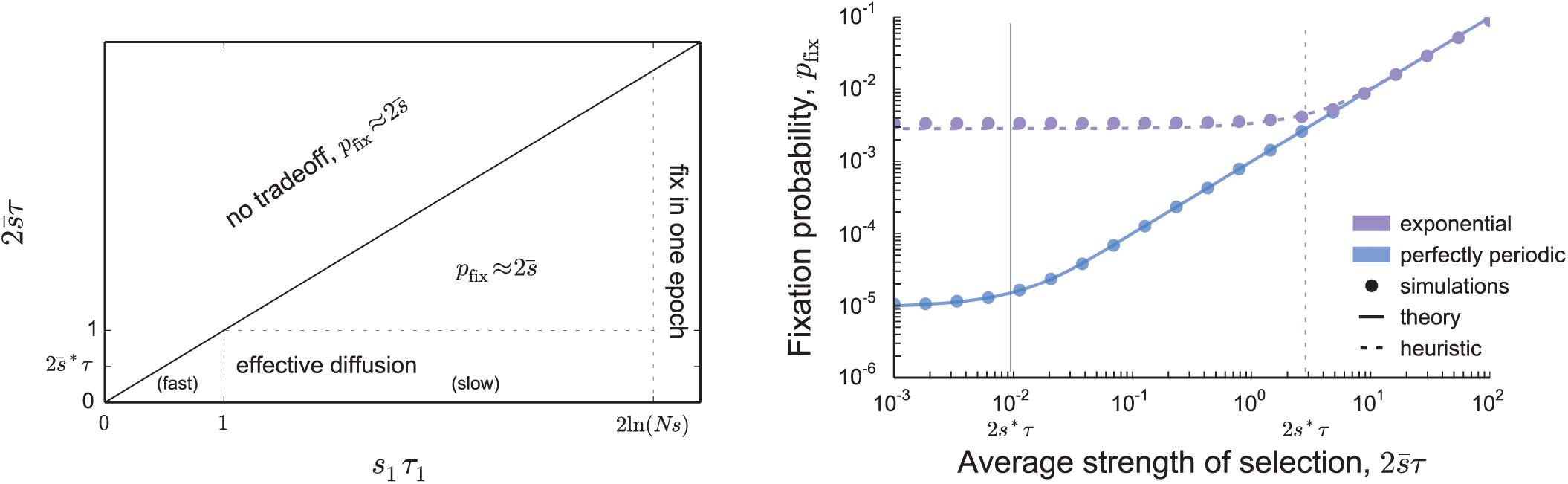
Left: Phase diagram showing the regime in which the average fitness effect is substantial, as a function of 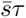 and *s*_1_*τ*_1_. Right: Comparison between the predictions obtained and Wright-Fisher simulations for perfectly periodic environments (*δτ* = 0, purple) and for exponentially distributed epoch lengths (*δτ* = *τ*, blue) (*N* = 10^6^, *s*_1_ = 10^−2^, *s*_1_*τ* = 10, 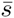 varied from 10^−6^ to 10^−1^).

In addition, when *δτ* ≳ *τ*, the precise shape of the epoch length distribution starts to play a larger role, and the mean and variance may not adequately capture the behavior. To see how these effects can become important for even larger *δτ*, we can consider the fixation probability under a gamma distribution of epoch lengths with *δτ* ≫ τ:

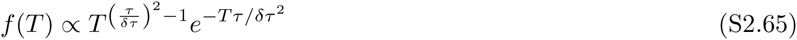

In this case, the distribution of epoch lengths resembles a power-law distribution with a median near τ and occasional fluctuations that are cut off at *δτ*^2^/*τ* ≫ *τ*. However, unlike the cases above, the duration of the very first epoch (i.e., conditional on a mutation arising) is dramatically different from that of a typical epoch, since it satisfies

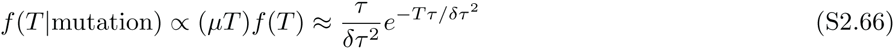

and has a typical scale much larger than τ. Provided that 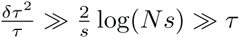, the mutation is virtually guaranteed to fix in its first good epoch, even though it would rarely expect to see an epoch of that length again. Thus, in the limit that *δτ/τ* → ∞ we again converge to the fix-in-one-epoch limit *p*_fix_ ≈ s. However, the next-order correction is much more difficult to obtain in this case, since the standing variation from an incomplete sweep will no longer be completely purged in the subsequent deleterious epoch. Rather, the dynamics of the mutation resembles a mixture of the diffusive dynamics in Section I with occasional jumps that can potentially drive the allele to fixation or extinction (similar to the generalized diffusion models studied in Ref [41]). A detailed analysis of this regime is beyond the scope of the present paper, and remains an interesting avenue for future work.

### III. RELATION TO PREVIOUS WORK

In the present work, we have focused on a diffusion model (Eq. 1) for the frequency of an allele in a fluctuating environment. This model bears many similarities to those employed in earlier studies of time-varying selection pressures, but it differs from these earlier models in several key ways. It is therefore useful to briefly review this earlier literature, so that we may comment on the major differences that arise.

The earliest attempts to model the effects of fluctuating selection pressures were largely focused on infinite-population models in which the selection coefficient is resampled from some fixed distribution in every generation, and the log-frequency of the allele undergoes a discrete random walk [12–14, 17–21, 26, 31]. In our present terminology, this is effectively a model of pure seasonal drift. Yet we have seen that while seasonal drift shares the dispersive nature of genetic drift, its multiplicative nature ensures that it can never completely drive an allele to fixation or extinction. Rather, the allele frequencies start to accumulate near *x* = 0 or *x* = 1, an effect which has been termed quasi-fixation [26]. However, just a small amount of genetic drift (or equivalently, a population size just slightly less than infinity) is sufficient to eliminate this pathological behavior [26, 31].

More general diffusion models were later proposed to account for the joint effects of seasonal variation and genetic drift [27–31]. Like their earlier counterparts above, these models assumed that selection pressures were resampled every generation, so that the standard derivation of the diffusion equation could still be applied [35]. However, due to this assumption of rapid and uncorrelated environmental change, these studies found that the effects of fluctuations are relevant only when the *variance* in the selection pressure is large compared to the other selection pressures in the population. This requires a modified version of the standard diffusion limit to account for the fact that 𝒪(*s*^2^) terms are no longer negligible, which raises a host of additional issues that can usually be ignored when *s* ≪ 1. In particular, the precise details of the birth-death process start to become important, as do differences in the definition of the selection coefficient, and whether time averages should be carried out using the geometric mean or the arithmetic mean [15, 21]. Somewhat more interestingly, these large-*s* effects can lead to an emergent form of overdominance, even for otherwise semi-dominant alleles [9]. When this occurs, the boundaries of the diffusion process are no longer accessible, and the mutation can be maintained at intermediate frequencies for extended periods of time [42–45].

However, we stress that all of these effects are absent in the standard diffusion limit (i.e., *N*→ ∞ and *s* → 0 with *Ns* held fixed), which is employed throughout the present work. Given this assumption, our model bears the closest similarity to the one employed by [32, 33], who considered autocorrelated selection pressures for which 1 ≪ τ ≪ 1/*s*. By exploiting an unused degree of freedom in the diffusion timescale, these authors derived an effective diffusion process not unlike the one considered here, and were the first to show that environmental fluctuations can be relevant even in the standard diffusion limit, provided that the *integrated* autocovariance in *s*(*t*) is larger than 1/*N*.

